# Commentary on Pang et al. (2023) Nature

**DOI:** 10.1101/2023.10.06.561240

**Authors:** Kaustubh R. Patil, Kyesam Jung, Simon B. Eickhoff

## Abstract

Pang et al. (2023) observe that the geometric eigenmodes, derived from the shape of the cortical surface, are better at reconstructing patterns of both spontaneous and stimulus-evoked activity, when contrasted with three alternative connectome-based models including structural connectome derived eigenmodes. Based on this observation they propose that geometric eigenmodes offer a good model for explaining brain function, noting that “*wave dynamics offer a more accurate and parsimonious mechanistic account of macroscale, spontaneous cortical dynamics captured by fMRI*”. They then question the prevailing view that brain activity is “*localized to focal, spatially isolated clusters*” and it is driven by “*intricate patterns of anatomical connections*”. While the observation that geometric properties fit brain activity well is intriguing, we argue that accepting geometric eigenmodes as a model for brain function risks the logical fallacy of “affirming the consequent”. A representation that effectively describes the underlying geometry is inherently adept at fitting patterns within that geometric space; it does not necessarily shed light on mechanisms of the brain’s functional attributes. To this end, we provide two lines of empirical results: (a) Basic parcel-based representations, which capture localized structures, can reconstruct activity patterns as well. (b) Geometric eigenmodes demonstrate a high flexibility when fit to a range of manipulated patterns, which evokes the danger of overfitting. Based on those results, theoretical considerations, and previous data we argue that more consideration is needed regarding “parsimony, robustness and generality of geometric eigenmodes as a basis set for brain function”. While we recognize the potential role of the brain’s geometry in influencing its dynamics, assertions regarding its efficacy should be weighed against the performance of simpler models, an inherent risk of overfitting and anatomical evidence. Pang et al.^1^ put forth harmonic modes derived from the brain’s geometry as a previously underrecognized model to explain brain-wide dynamics. Their reconstruction framework relies on multiple linear regression to fit brain patterns using a basis set and then calculating Pearson’s correlation between original data and fitted data, both parcellated using an atlas with 180 parcels in each hemisphere^2^. In addition to the overarching challenge of accepting any model as an actual reflection of the real world^3^, we here provide empirical results and theoretical arguments that highlight the need for further consideration regarding geometric eigenmodes as a model of macroscale brain activity.

## Simple localized models can reconstruct activity patterns

None of the basis sets compared by Pang et al. explicitly model localized cortical activity. To this end, using the same reconstruction framework, we reconstructed brain activity patterns using a basis set derived from a widely used parcellation scheme^4^, which has been shown to converge well with histologically defined brain areas. This discrete and orthogonal basis set following classical definition of cortical areas encoding vertex membership outperformed the connectome and was comparable to the EDR eigenmodes (Fig 1a-b). When enhanced using the structural connectome it provided even better reconstruction than geometric eigenmodes, particularly when using smaller basis sets (50 modes in Fig 1a, Suppl. Fig 1). Similar observations were made when using parcels obtained by clustering the geodesic distance matrix (Suppl. Fig 2). These results suggest that “discrete, isolated and anatomically localized” clusters in fact reconstruct brain activity patterns without the need for harmonic information.

**Figure 1.**
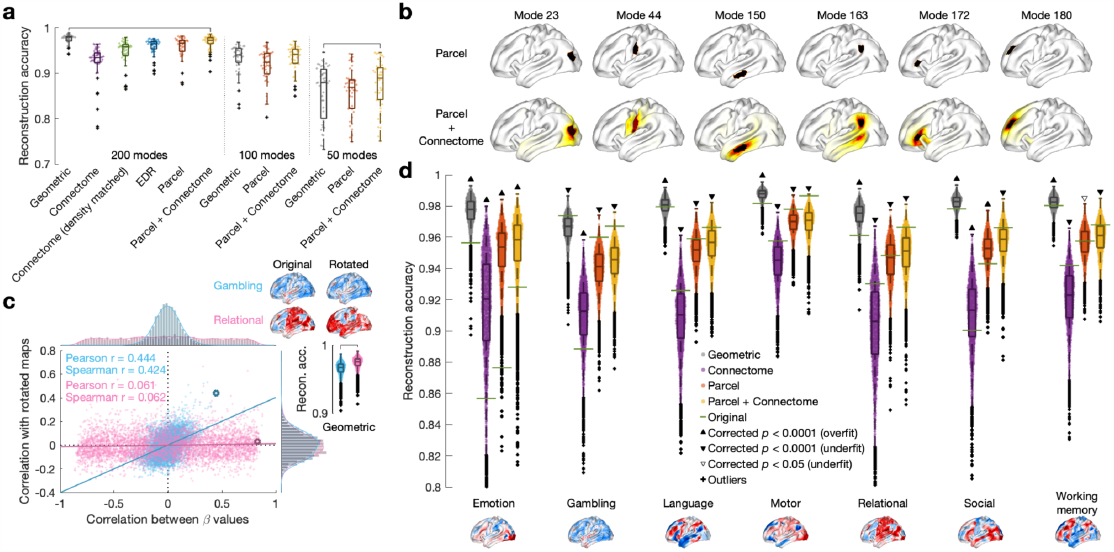
**a**, Reconstruction of 47 HCP task contrast maps using different modes at three different granularities. We used the Schaefer atlas of appropriate size to obtain the parcel modes. To obtain the Parcel+Connectome modes we averaged the structural connectome values for the vertices within a parcel and assigned the maximum value to the parcel vertices. The horizontal lines represent significant statistical differences using the Wilcoxon signed-rank (paired) two-tail test. We only compared geometric eigenmodes with Parcel+Connectome modes. EDR: exponential distance rule. **b**, Illustration of selected Parcel and Parcel+Connectome modes. **c**, Reconstruction of two of the HCP key task contrasts and their randomly spun versions (N=5000). The correlations were calculated between that of original data and spun data (the beta value of the first eigenmode was excluded). For the Gambling task we see a positive correlation between task map similarity and that of the corresponding beta values (spun data compared with original data, see the example maps on the top-right corner corresponding to the circled points in the scatter plot). For the Relational task no relationship was observed. **d**, Reconstruction of randomly spun data for the seven key HCP tasks. In all but one case (Gambling) the geometric eigenmodes fit the spun data significantly better (Wilcoxon signed-rank one-sample two-tailed test and Bonferroni correction) than the original data (green horizontal lines). The other modes generally tended to underfit the manipulated data compared to the original data.

## Geometric eigenmodes may affirm the consequent

Next, we contend that the impressive reconstruction accuracy of geometric eigenmodes may stem from their descriptive nature rather than mechanistic effects. If “wave dynamics unfolding on the geometry of the cortex” truly serves as a “generative mechanism for capturing complex properties of spatiotemporal brain activity”, one might reasonably expect similar activity patterns to emerge from the excitation of similar eigenmodes. Yet, in most tasks, this expectation does not hold and similar and dissimilar brain patterns are accurately fit by correlated beta values (Fig 1c, Suppl. Fig 3). In addition, geometric eigenmodes were more susceptible to fitting different types of randomly manipulated data (Fig 1d, Suppl. Fig 4, Suppl. Fig 5). This tendency may reflect the capacity of geometric eigenmodes to fit any patterns rather than (only) biologically plausible ones even when derived from non-brain shapes^5^. These findings raise questions about generative and explanatory capacity of geometric eigenmodes.

These empirical findings open the door to several theoretical and data-related considerations. The effectiveness of data-fitting depends on how well the basis set describes the underlying vector space. The Laplace-Beltrami operator (LBO) was devised to accurately and concisely capture the morphology of a shape^6,7^. Given this descriptive nature, LBO-based geometric eigenmodes are expected to represent the shape and each vertex better (Suppl. Fig 6) and hence accurately fit patterns on this shape. This may lead to the risk of “affirming the consequent”, i.e., geometric descriptors can accurately fit geometric patterns. This argument is reinforced by a straightforward demonstration wherein parcel-modes derived from the same 180 parcels used for evaluation result in nearly perfect reconstruction (Suppl. Fig 2c). A high reconstruction accuracy thus reflects the descriptiveness of the basis set and does not necessarily imply a mechanistic explanation.

The utility of a model is contingent on its ability to make testable predictions rather than the mere capacity to fit/accomodate a large volume of data^8^. Conversely, as noted above, using a sufficiently fine-grained and well-defined basis set may become susceptible to overfitting. It remains unclear whether the geometric eigenmodes as a model of brain function can effectively make forward-looking predictions regarding brain activity patterns that have not yet been observed. Consequently, the generative aspect of this model is yet unclear.

Interestingly, Pang et al. posit geometric eigenmodes are a more parsimonious model, which should be simpler, all else being equal. Asserting parsimony would require explicitly comparing the simplicity of the basis sets because they are not nested, rather than merely comparing their cardinality. This admittedly very challenging task remains unaddressed. In turn, one could reasonably argue that parcel-wise modes are in fact a simpler representation and hence provide a more parsimonious model. However, we refrain from making this assertion pending concerns about overfitting and unknown predictive ability before considering them as a valid model^3,8^. Similar considerations arise when comparing the two dynamical simulation models which should not be compared just based on number free parameters as they are of completely different forms. Furthermore, the comparative evidence of dynamical models is rather limited given that the mass model is sensitive to parameters including the parcellation scheme^9^.

Additionally, the faithfulness of diffusion data in representing the structural connectome is a longstanding issue in the field. While Pang et al. adhered to standard processing procedures, inherent caveats still exist. In theory, a connectome provides a comprehensive depiction of axonal fibers. In practice, however, common MRI protocols–including the HCP protocol–lack the spatial resolution to capture intracortical axon collaterals and U-fibres^10,11^ that make up more than 95% of all axonal fibres^12^. Their short length also makes them crucial for capturing geometric properties. This is an important caveat to acknowledge when using connectome data and comparing it to other modalities together with additional considerations^13^.

Finally, we note that while geometric eigenmodes represent an interesting perspective on brain organization, it seems very hard to reconcile them with the well established topographic organization of the cerebral cortex^14^. This discrepancy is particularly challenging when considering that histological patterns such as cyto- and myeloarchitectonic borders, which can be directly observed through microscopy^15^ rather than being inferred through modeling and analysis, may be considered to have a higher epistemic value. In other words, we would argue that any model-based proposal for brain organization should be able to accommodate directly observable evidence on cortical topography.

In summary, the *sine qua non* of the geometric (and possibly other) eigenmodes may stem more from mathematical underpinnings than biological factors. Further investigations are needed to truly grasp the nature and behavior of such models–both to understand why they fit the data so well and to realize their full potential by devising strategies to control potential overfitting.

## Disclaimer

The presented opinions are those solely of the authors and do not necessarily represent the opinions of their employers.

## Acknowledgements

This work was partly supported by the Helmholtz Imaging Platform project NimRLS.

We thank Prof. Juergen Dukart and Dr. Casey Pacquola, and Dr. Oleksandr Popovych for insightful comments on an early version of this commentary.

## Supplementary Material

The data provided by Pang et al. were downloaded from the two online resources: https://github.com/NSBLab/BrainEigenmodes https://osf.io/xczmp

Parcel modes: These were defined as a binary matrix that encoded each parcel as a mode, i.e. column of the matrix, with the vertices assigned to that parcel assigned a 1 and rest of the vertices as 0.

Parcel+Connectome modes: These were modifications of the parcel modes where for each parcel the average structural connectivity was also considered. To emphasize the parcel’s nature, the max value was assigned to each vertex assigned to that parcel.

Reconstruction: Unless explicitly specified we performed reconstruction on the brain activity patterns averaged across the 255 participants as provided by Pang et al. A multiple linear regression model was fitted and the fitted values and the real values were compared using Pearson’s correlation after averaging over 180 parcellations from the Glasser atlas. We used only the left hemisphere data.

Random manipulations: Averaged task maps were randomly manipulated either using spin permutation or Moran randomization using the BrainSpace toolbox.

**Suppl. Figure 1.**
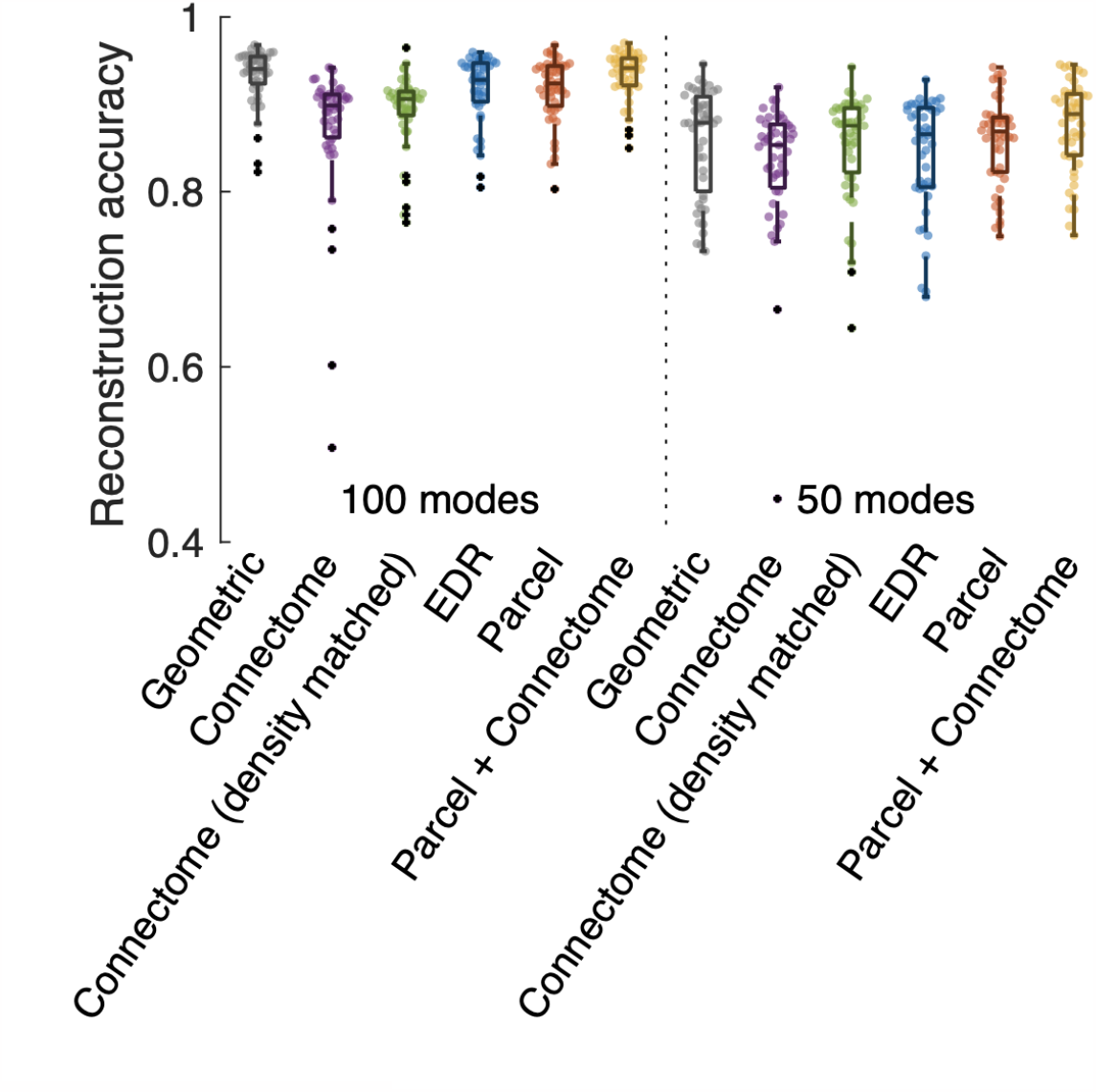
Reconstruction accuracy of 47 HCP task contrast maps using 100 and 50 modes in using different basis sets. The parcels in the left hemisphere of the Schaefer atlas of 100 and 50 parcels (200 and 100 parcels for both hemispheres, respectively) were used for the ‘Parcel’ and ‘Parcel + Connectome’ basis conditions. EDR: exponential distance rule.

**Suppl. Figure 2.**
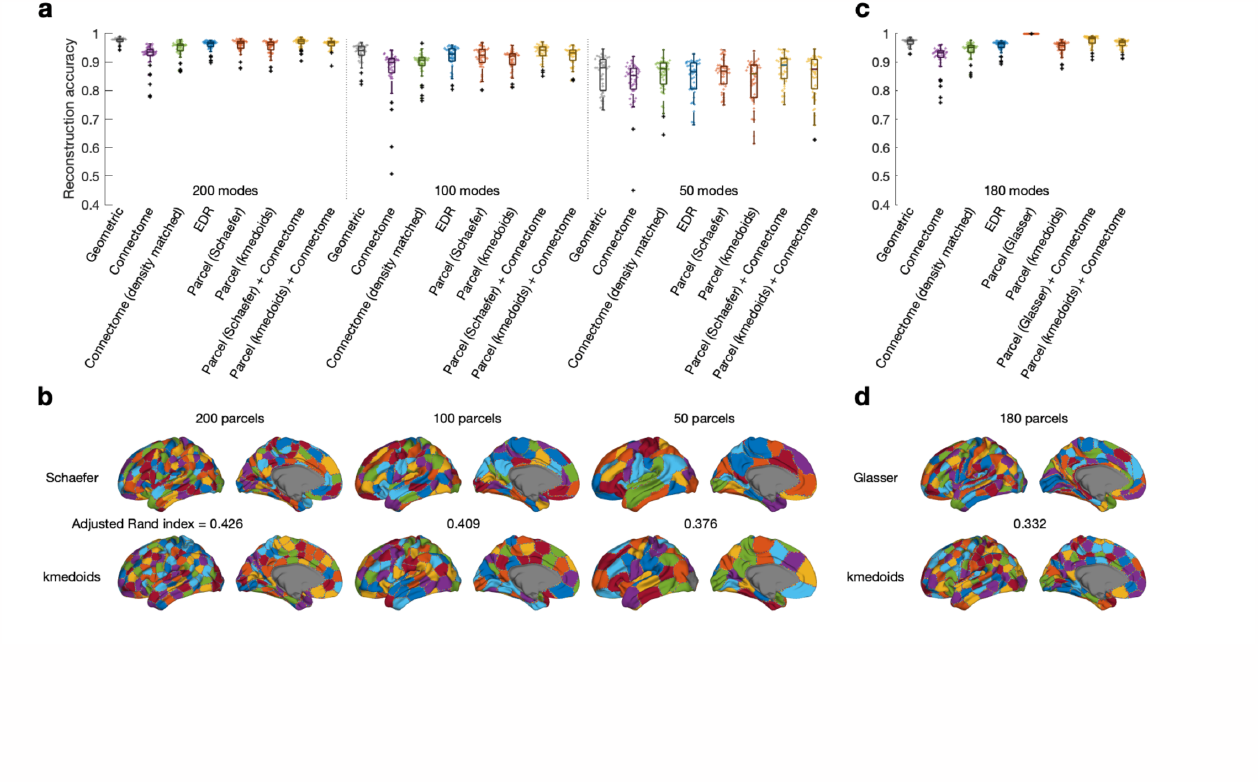
Comparison of existing parcellation schemes and that derived from clustering of the geodesic distances. The vertex-wise geodesic distance matrix (without the medial wall) was clustered using the k-medoids algorithm into the required number of clusters. These clusters were used for deriving the Parcel and Parcel+Connectome modes and subsequently for reconstruction of the 47 HCP task maps for the Schaefer parcellations with 400 (200 modes), 200 (100 modes), and 100 (50 modes) parcels (**a**) and the Glasser atlas with 360 parcels (180 modes, **c**) on the left hemisphere The k-medoids derived modes generally showed slightly lower reconstruction accuracy than the known parcellations though still comparable to that of connectome-based eigenmodes. **b, d**, The k-medoids derived clusters showed a moderate level of match based on the adjusted Rand index with the corresponding Schaefer parcellations and Glasser parcellation. The parcellations also showed moderate similarity to the 180 parcels from the Glasser atlas with 0.332: 200 parcels 0.338 and 0.318, 100 parcels 0.365 and 0.337, and 50 parcels 0.294 and 0.263 (Schaefer and k-medoids, respectively). EDR: exponential distance rule.

**Suppl. Figure 3.**
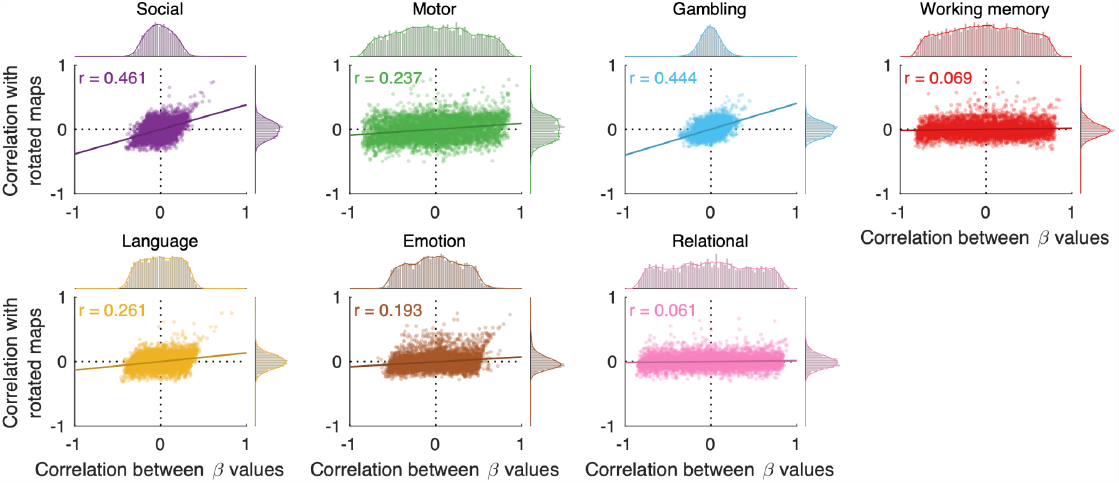
The seven HCP key task contrasts (averaged across 255 subjects) and their randomly spun versions (N=5000) were reconstructed using 200 geometric eigenmodes. Each scatter plot shows correlation between task map similarity (y-axis) and that of the corresponding beta values (x-axis, the first beta value corresponding to the 0 eigenmode was discarded). The r-value in the top-left corner is Pearson’s correlation coefficient of the two corresponding correlation values. For a consistent model it is expected to observe a positive correlation indicating that similar beta values are used to reconstruct similar brain maps.

**Suppl. Figure 4.**
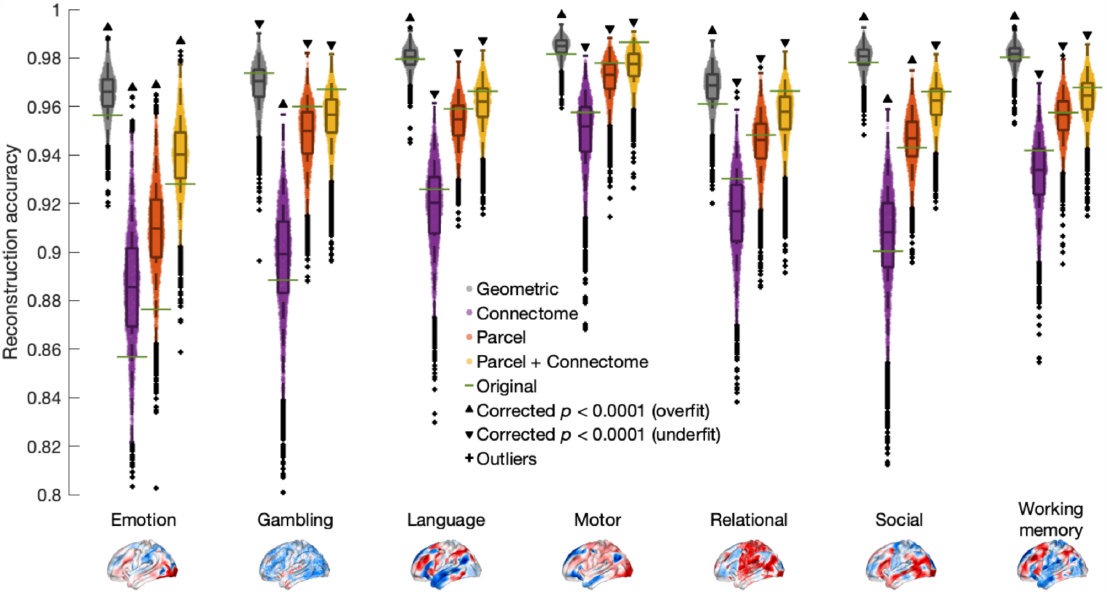
Reconstruction of manipulated data generated by adding randomly spun data to original maps for the seven key HCP tasks. In all but one case (Gambling) the geometric eigenmodes fit the manipulated data significantly better (Wilcoxon signed-rank one-sample two-tailed test and Bonferroni corrected statistics) than the original data (green horizontal lines). The other modes generally tended to underfit the manipulated data compared to the original data.

**Suppl. Figure 5.**
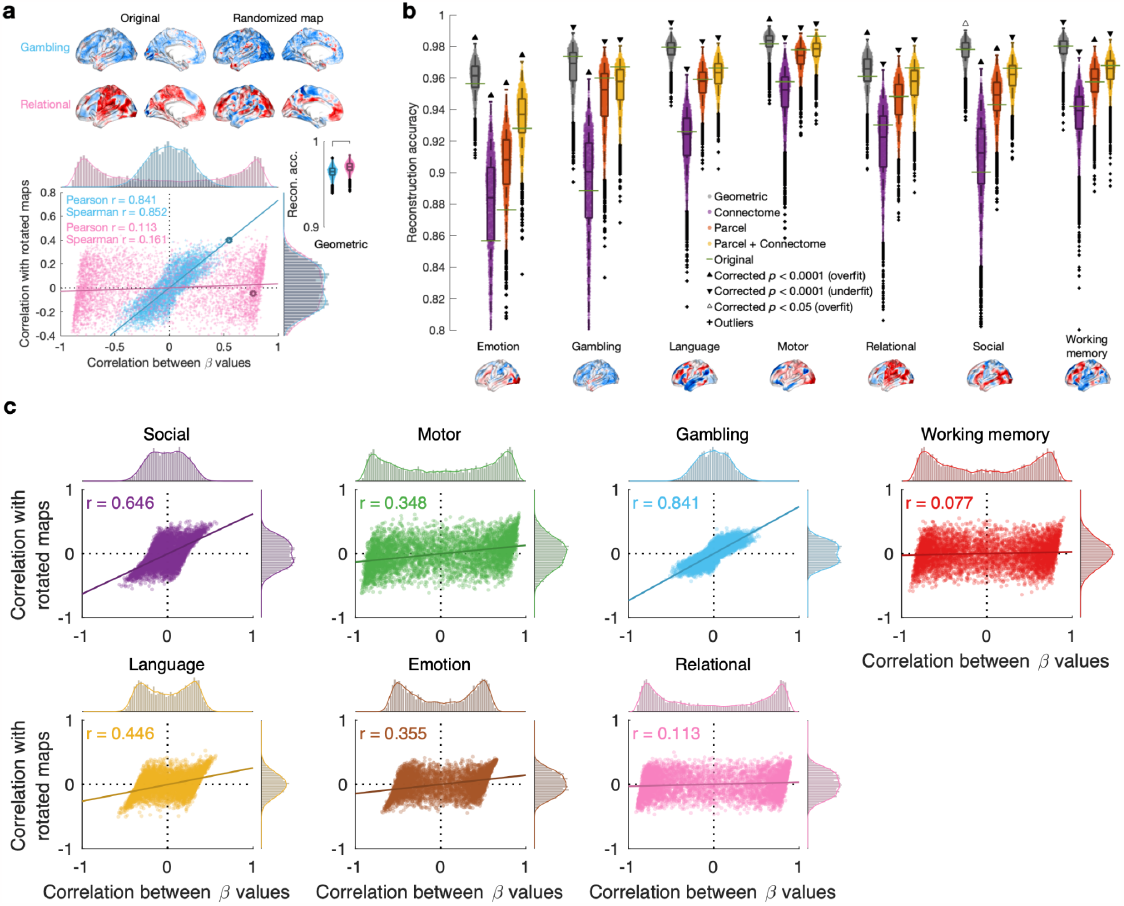
**a**, Reconstruction of two of the HCP key task contrasts and their Moran randomized maps (N=5000). The correlations were calculated between that of original data and random data as well as the corresponding beta values obtained for reconstruction (the beta value of the first eigenmode was excluded). For the Gambling task we see a positive correlation between task map similarity and that of the corresponding beta values (randomized data compared with original data, see the example maps on the top rows corresponding to the circled points in the scatter plot). For the Relational task no relationship was observed. **b**, Reconstruction of randomized data (Moran) for the seven key HCP tasks. For some key tasks the geometric eigenmodes fit the randomized data significantly better (Wilcoxon signed-rank one-sample two-tailed test and Bonferroni correction) than the original data (green horizontal lines). The Parcel (except Emotion and Social) and Parcel+Connectome (except Emotion) modes tended to underfit the manipulated data compared to the original data. **c**, Scatter-plots for all seven HCP key tasks similar to the one shown in the panel a.

**Suppl. Figure 6.**
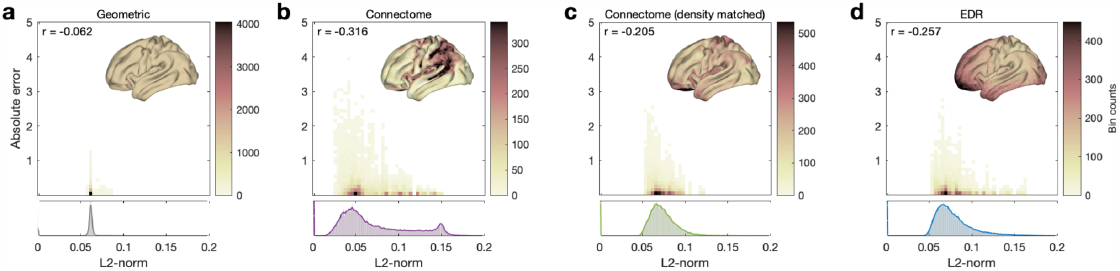
Heatmaps of L2-norm versus absolute reconstruction error values for each of the four eigenmodes. The reconstruction errors based on 200 eigenmodes of all 47 HCP tasks were calculated as absolute difference between task contrasts and reconstructed maps in averaged values using the Glasser atlas (180 parcels on the left hemisphere). The values shown in the top-left corner indicate Pearson’s correlation between the L2-norm and the absolute error. The cortical maps show the patterns of L2-norm on the cortical surface. The histograms at the bottom show distributions of L2-norm values (scaled). The geometric eigenmodes cover all the vertices in a more uniform manner while the connectome eigenmodes do not. This coverage (i.e. L2-norm) negatively correlated with the reconstruction error for all three non-geometric eigenmodes. This suggests that some eigenmodes lack coverage of the vector space to be able to reconstruct a signal of interest. EDR: exponential distance rule.

